# Convolutional Neural Network Analysis of Social Novelty Preference using DeepLabCut

**DOI:** 10.1101/736983

**Authors:** Nicholas B. Worley, Anthony Djerdjaj, John P. Christianson

## Abstract

The description and quantification of social behavior in laboratory rodents is central to basic and translational research. Conventional ethological approaches to social behavior are fraught with challenges including bias, significant human effort and temporal accuracy. Here we show proof of principle that machine learning can be applied to laboratory tests of social decision making. Rats underwent social novelty preference tests which were scored both by hand and again by a convolutional neural network generated in the DeepLabCut computer vision package of Mathis and colleagues. The CNN generated temporally (30Hz) and locally (<5pixels) accurate identification of rat nose, eye and ear positions which were then used to compute social interaction and topography heat maps. In sum, hand- and computer-scoring were strongly correlated, and each identified significant preferences to interact with novel conspecifics which sets the stage for applying DeepLabCut analysis to other types of social interaction in the future.

## Introduction

Abnormalities in social cognition underlie autism spectrum disorder (ASD) and numerous other psychiatric conditions (Decety, Skelly, & Kiehl, 2013; Thioux & Keysers, 2010). A major part of basic and translational research into the pathophysiology of ASD has involved studies of rodent (mouse and rat) social behaviors and how various factors (genetic, environmental, and drug treatment) alter the patterns of social behavior that can be observed in laboratory rodents (Pasciuto et al., 2015). Rats and mice are generally social animals that engage in exploratory behaviors with social partners (often measured as “sociability,” Nadler et al., 2004). When given the choice between familiar or novel conspecifics, investigation is typically directed more to the novel target (Barefoot, Aspey, & Olson, 1975). Importantly, genetic variability (i.e. between strains of rodents), mutations (e.g. SNPs or knockouts relevant to ASD), and drug treatments have all been found to influence the rodent’s tendency for sociability and novelty which make rodent social behavior studies useful for preclinical research (Kazdoba et al., 2015).

The majority of investigations into social behavior in laboratory rodents have taken an ethological approach in which trained observers time or count the number and duration of episodes of specific types of behaviors as they occur over time. For example, in a simple social encounter between 2 subjects, an observer could quantify social grooming, sniffing, pinning, chasing or mating behaviors. The generation of ethograms is typically aided by computer timing programs and video which allow slowing of video and multiple observer scoring for validation and inter-rater-reliability estimations. This method is very effective and has resulted in scores of publications. However manual scoring of social behavior videos is fraught with some serious challenges (Anderson & Perona, 2014). First, any human observer is at risk of making biased observations. This can reduce the rigor and reproducibility of experimental results. Second, human timing is limited in temporal accuracy in so far as the beginnings and endings of defined behaviors will be offset by the reaction time of the observer which is likely to be highly variable and can make synchronization and integration of social behavior with physiological recordings, such as single unit electrophysiology very difficult. Third, human scoring requires a significant amount of experimenter hours.

Advances in machine vision for object recognition set the stage for a possible to solution to all 3 of the problems described above. DeepLabCut is one such development which utilizes a supervised learning approach in which a human expert trains a convolutional neural network (CNN) to track a set of patterns or features in the frames of video (Mathis et al., 2018). DeepLabCut is remarkably useful for training a CNN to identify the poses of animals in video data without the need of specialized marking (such as reflectors). The goal of this report is to document the application of DeepLabCut in training a CNN to identify rat locations during a social novelty preference test in which an experimental rat is given the choice to investigate either an unfamiliar conspecific or a familiar, cagemate in an arena. This is a well-established phenomenon in which the experimental rat is expected to spend much more time investigating the unfamiliar conspecific than its cagemate (Smith, Wilkins, Mogavero, & Veenema, 2015).

DeepLabCut was applied to 2 experiments. In the first, male rats were given social novelty preference tests in which the behavior of the test rat was quantified by either a trained observer or DeepLabCut over a 5-minute period. The experimental rat was able to freely explore an arena which contained the familiar and unfamiliar target rats in chambers on either end. The second experiment was, in principle, the same experiment except that the rats involved received an experimental treatment, described elsewhere (Rogers-Carter, Djerdjaj, Gribbons, Varela, & Christianson, 2019). The second experiment was again analyzed by a human observer and by DeepLabCut using the deep neural network trained on Experiment 1 videos in order to test the generalizability of learning algorithm.

## Materials and Methods

### Rats

The complete description of the procedures involving rats are published (Rogers-Carter et al., 2019). Briefly, male Sprague-Dawley rats were obtained from Charles River Laboratories (Wilmington, MA) and housed in same-sex groups of 2-4 with free access to food and water on a 12hr light/dark cycle. Experimental rats were housed and tested with same-sex conspecifics. Familiarity was established by cohousing experimental rats with conspecifics. Familiar group cages housed 2 experimental rats and 2 conspecifics; 1 of the conspecifics was used as the naive and the other as the stressed target for the SAP tests of the 2 experimental cagemates. Each conspecific cage-mate pair was used for 2 SAP tests. The rats used for targets in SAP tests with unfamiliar conspecifics were housed separately in groups of 4. Rats were housed in these groups for 7 days prior to testing. Behavioral tests were conducted within the first 4hr of the light phase and all procedures involving rats were conducted in accordance with the *Guide for the Care and Use of Laboratory Animals* and approved by the Boston College Institutional Animal Care and Use Committee.

### Social novelty preference test

The social novelty preference test allows for the quantification of social exploration of an experimental rat directed toward 2 target conspecifics, 1 familiar (cagemate) and one novel. The social novelty preference test began with 2 days of habituation: experimental rats were acclimated to the test area, a clear plastic cage (50 x 40 x 20cm, L x W x H) with a wire lid, for 60 min and then presented 2 empty restraint chambers (day 1) or 2 naive conspecifics (day 2) housed in the restraint chambers. Restraint chambers were clear plastic enclosures (18 x 21 x 10 cm, L x W x H; see Rogers-Carter, Djerdjaj, Culp, Elbaz, & Christianson, 2018) which allow for social exploration through acrylic rods spaced 1 cm center-to-center. To assess social novelty preference on day 3 the experimental rat was presented 2 conspecifics, 1 familiar cagemate and one novel. Social novelty preference tests were recorded on digital video. Human scored social exploration was defined as any time the experimental rat made physical contact with a conspecific and was quantified for both the naive and stressed targets. Social exploration was quantified during live testing and again from video by an observer who was completely blind to experimental conditions. Behavioral videos were collected with a 8.9 megapixel Sony Handicam (model number HDR-PJ430). Videos were recorded at a resolution of 1440×1080, a bitrate of 5358kbps, frame rate of 30 frames per second and converted to a final resolution of 720×480, a bitrate of 276kbps at 30fps using Handbrake (Version 1.2.2).

### Computer and DeepLabCut Installation

Training, evaluation, and interaction zone analysis were performed on a custom-built computer containing and Intel Core i5-4690K CPU, 3.50Ghz, 8.00 GB RAM, EVGA GeForce RTX 2080 graphics processing unit 8GB, and running Windows 7 64-bit operating system. Python was installed through Anaconda 2.2 and DeepLabCut 2.0.7 was installed in an anaconda environment with Python 3.6.8 and Tensorflow 1.13.1 (see Appendix 1 for full list of environment packages and versions).

### DeepLabCut Network Training

480 frames were extracted uniformly from videos (12 frames per video) of 40 distinct social novelty preference tests and manually labeled for nose, eyes, ears (Figure 1). The data were then split randomly into training set of 456 frames (95% of extracted frames) and test set of 24 frames (5% of extracted frames). A CNN was trained using the 456 frames of the training set from for 1,030,000 iterations. Evaluation of labeling accuracy was achieved by comparing the labels acquired from the CNN on the test set with manual labels. The model was then used to evaluate all frames in each of the 40 videos used for training. The resulting *x* and *y* coordinates corresponding to the nose position within each frame were used to determine time spent in the interaction zone. The interaction zones were defined by a polygon containing the four points at each corner of the center-facing wall of the conspecific-containing corrals and padded by 25 pixels toward the center of the arena for a total width of ~3cm. The sum of the total number of frames where the nose position was within the boundary was multiplied by the frame rate (30 fps) to achieve total seconds spent in the interaction zone. Heatmaps were constructed using numpy as follows. The center items (2×2) of a 10×10 array containing zeros was set to a value of 200 and a gaussian blur was applied to produce a “dot” to be added to the heatmap. The heatmap was produced by first initializing a 720×480 array containing the value of 255 for each element. The dot was then subtracted from the array using the nose coordinate labeled by the CNN. Each array was then threshold to a minimum value of 0, arrays from behavioral tests with the familiar conspecific on the left were then flipped about the *x*-axis, and the average of all arrays was plotted using Matplotlib with a red-blue colormap, with red corresponding more time spent in a given location.

**Figure 1.**
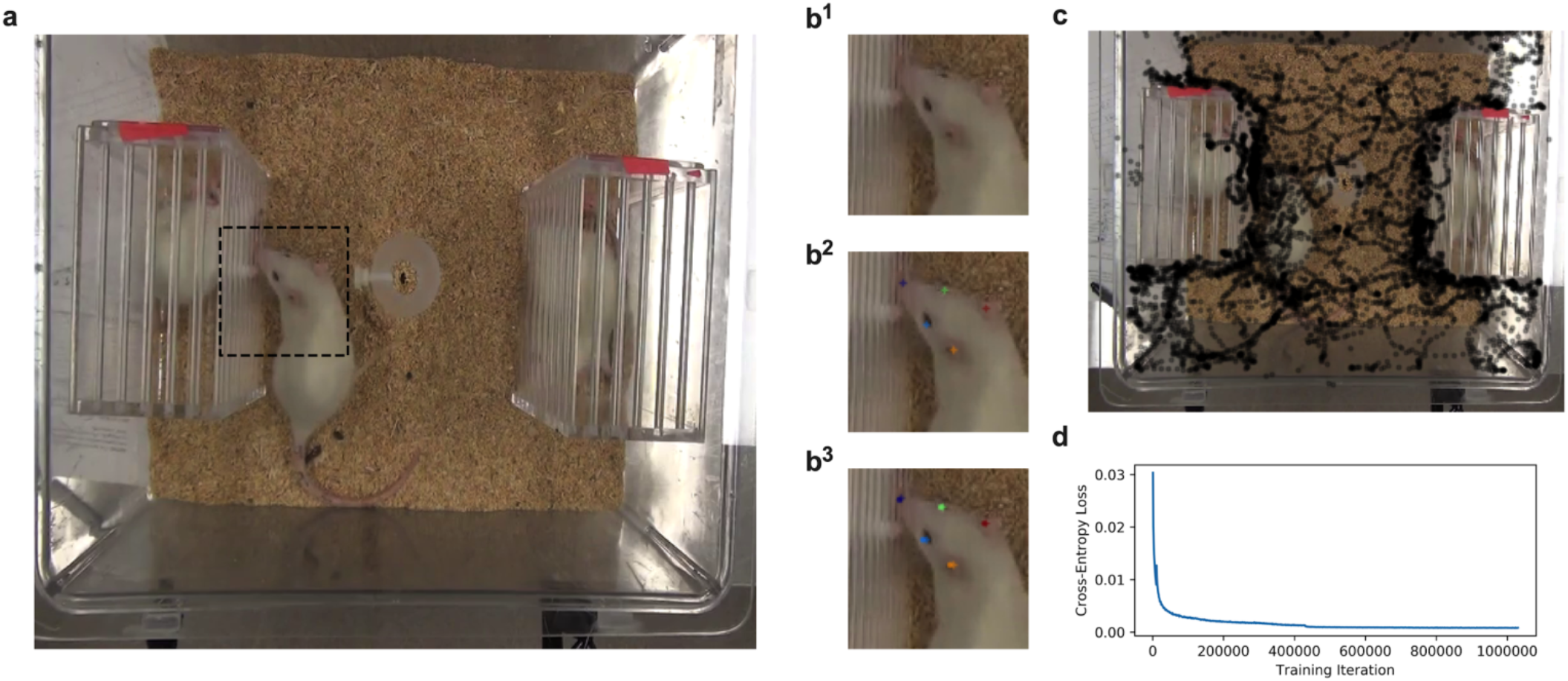
DeepLabCut Training. **a)** Example of a video frame used for pose labeling and convoluted neural network (CNN) training. The experimental rat was free to explore the arena. Unfamiliar or Familiar target rats are contained within plastic chambers at either side of the arena. **b^1^)** Unlabeled image. **b^2^)** Nose, eye, and ear position markers placed by human expert to train the CNN. **b^3^)** Nose, eye, and ear positions determined by DeepLabCut. **c)** Topography of rat nose position across time in the test. **d)** CrossEntropy Loss decreased over iterations while training the CNN.

## Results

Training of the CNN through 1,030,000 iterations took 17h and 38m. Cross-entropy loss continued to decrease throughout the training process (Figure 1c) and the final model achieved an average error of 1.56 pixels on training images and average error of 5.39 pixels on test images.

Hand-scoring of the social novelty preference test revealed that the rats spent more time socially interacting with the novel conspecific than with the familiar conspecific (t_9_=3.703, p<0.01; Figure 2a). The trained CNN was then used to label all frames across the 40 videos, and these labels were used to determine time spent in the interaction zone (see Supplemental Video). Analysis of machine-labeled nose positions revealed that rats spent more time in the interaction zone of the novel conspecific than the familiar conspecific (t_9_=3.602, p<0.01; Figure 2b). Moreover, machine scoring was strongly correlated with handscoring (Pearson r=0.792, p<0.0001; Figure 2c).

**Figure 2.**
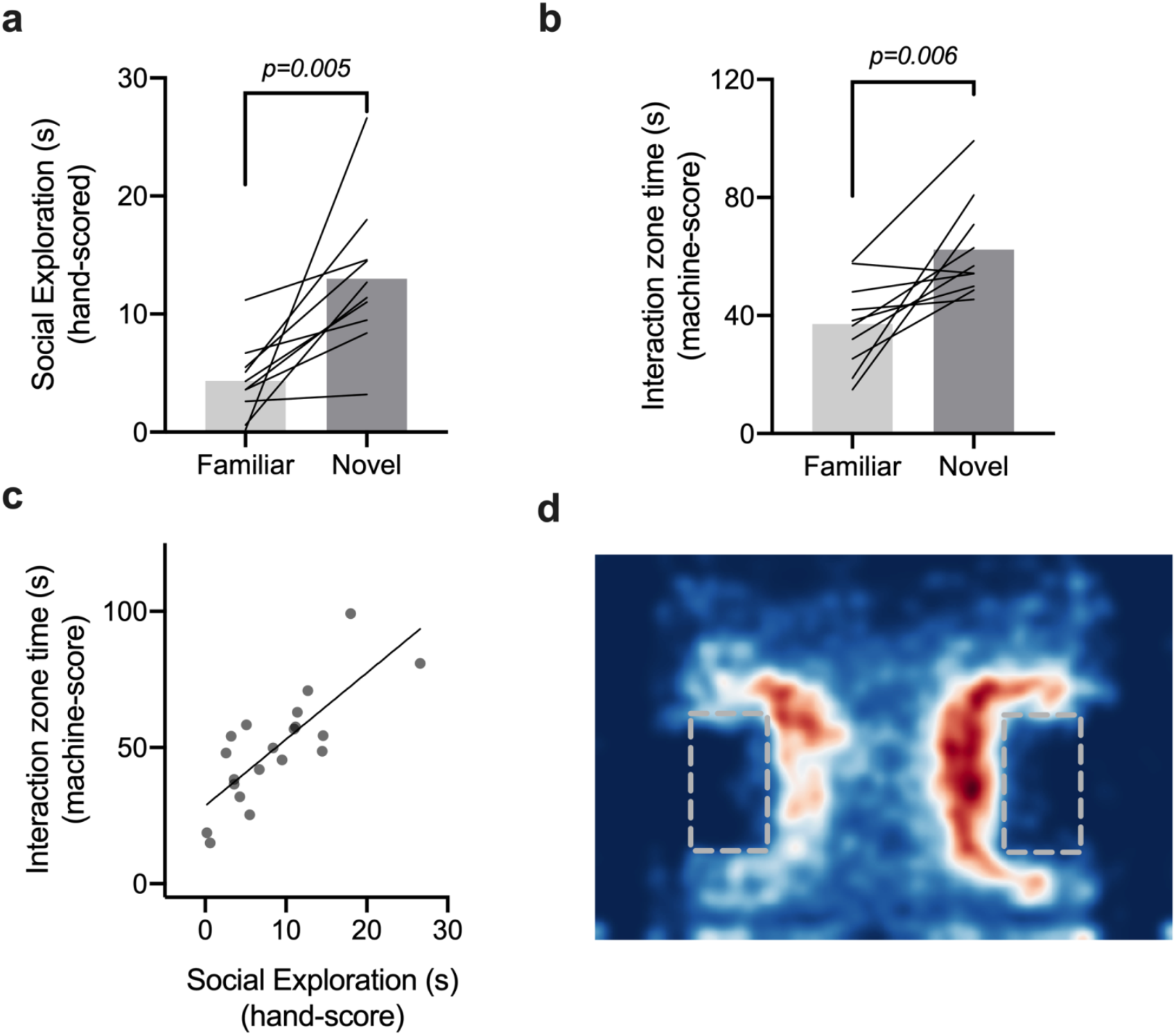
Time spent in the interaction zone of the familiar or novel conspecifics chamber determined by hand and by DeepLabCut computed nose position over 5 minutes. **a)** Time in seconds spent interacting with a familiar or novel conspecific, measured by a human observer. Rats spent significantly more time interacting with the Novel/unfamiliar subjects [paired samples t9 = 3.703, p = 0.005]. **b)** Time in seconds the test rat nose was found within the interaction zone as determined by DeepLabCut. The interaction zone was determined as a polygon drawn over the opening of the target chambers with 25 pixel padding. Noses tended to be found in the interaction zone of the novel/unfamiliar conspecific [paired samples t9 = 3.60, p = 0.006]. **c)** Correlation of hand scores and machine scores, [r = 0.79, p < 0.001]. **d)** Red-blue heatmap demonstrating relative time spent in the arena during the test. Red indicates more time spent in a given location while blue indicate less time. Gray dashed rectangles indicate the relative locations of the chambers containing the familiar conspecific (left) and the novel conspecific (right).

To determine the generalizability of the CNN, the trained CNN was used to analyze videos from another experiment with the same social novelty preference behavioral testing paradigm. Here, hand-scoring of the social novelty preference test revealed that the rats spent more time socially interacting with the novel conspecific than with the familiar conspecific (F(1, 6) = 19.12, p<0.01) regardless of drug treatment (F(1, 6) = 0.036, p=0.85; Figure 3a). Similarly, analysis of machine-labeled nose positions revealed that rats spent more time in the interaction zone of the novel conspecific than the familiar conspecific (F(1,6)=35.21, p<0.001; Figure 3b) regardless of drug treatment (F(1,6)=0.213, p=0.66; Figure 3b). Again, machine scoring was strongly correlated with hand-scoring (Pearson r=0.667, p<0.0001).

## Discussion

Here we demonstrate the use of DeepLabCut to quantify and visualize the behavior of rats in a laboratory social decision-making task. Behaviorally, we found that experimental rats choose to spend more time investigating a chamber containing a novel, unfamiliar conspecific compared to a familiar cagemate. This finding is consistent with a large literature on social novelty (Barefoot et al., 1975; Smith et al., 2015). Both hand scoring interaction time and the CNN scoring detected the same pattern which inspires confidence that more elaborate applications of DeepLabCut in similar settings will be useful. Nose position in the interaction zone captured 2-3 times more time in behavior than the strict hand scoring (only timing the direct social exploration of the target). The *x, y* coordinates provided enabled the generation of heat maps which help illustrate the pattern of behavior in the group.

**Figure 3.**
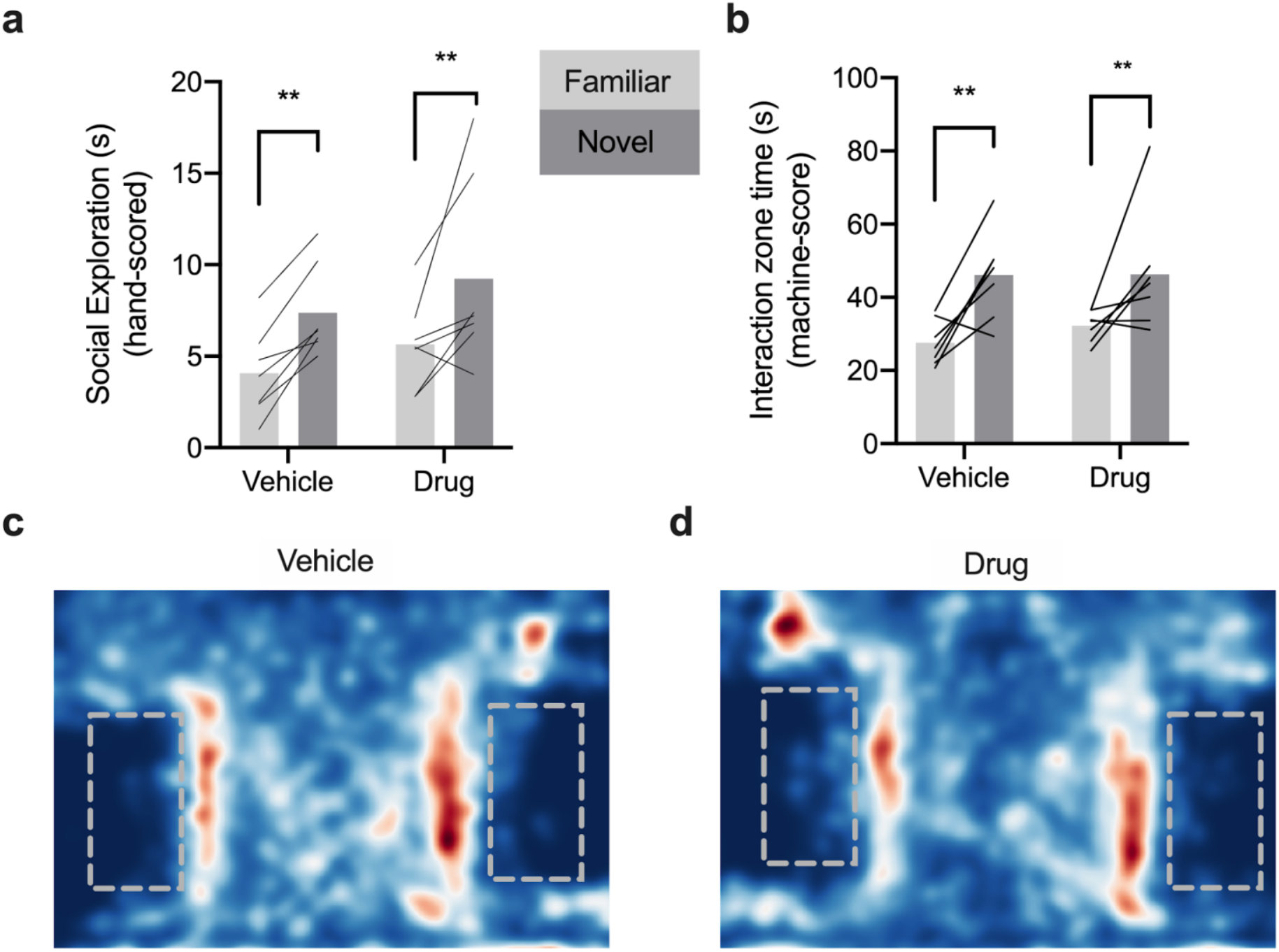
DeepLabCut performance on untrained video examples from experiment 2. **a)** Human scoring of social interaction shows that rats spent more time interacting with the novel conspecific under both vehicle and drug conditions as evinced by a main effect of familiarity [F1,6 = 19.12, p = 0.005]. **b)** The CNN that was trained on videos from Experiment 1 was used to process experiment 2 videos and nose time in the interaction zone was determined as in Figure 2. Rats spent more time in the interaction zone as evinced by a main effect of familiarity [F_1,6_ = 35.12, p = 0.001]. **c-d)** Heat maps indicating sum location of rat nose across all analyzed videos (n = 7 per condition). Gray dashes indicate the relative location of conspecific chambers, red indicates greater time and blue indicates less time. Data are aligned so that familiar conspecifics were located in the left chamber and novel/unfamiliar conspecifics were in the right chamber.

The benefits of computer based ethology are numerous (Anderson & Perona, 2014). A major goal of contemporary neuroscience research is to integrate correlates of neural activity, such as the firing of individual or groups of neurons, with specific behavioral or cognitive functions. A typical action potential is only a few ms long, faster than the human visual system can sense, appraise, and label the behavior of an animal (on the order of 100s of ms). Although this can be overcome with high speed video cameras, the task of identifying the start and end of specific behavioral bouts is often deemed too laborious to have value. Here, using a conventional digital video camera we are able to get timestamps of specific rat postures at a rate of 30Hz (~33ms) which opens the way for integration of posture data with neural recordings (although these may come at higher sampling rates). Another value of computer-based ethology would be improved statistical power for hypothesis testing. To explore this possibility, we compared the observed variance accounted for (effect size, *R^2^*) between the hand and computer scored ANOVAs. In experiment 1, *R^2^_hand_ = 0.60 and *R^2^_computer_* = 0.59 while in experiment 2, R^2^_hand_* = 0.22 and *R^2^_computer_* = 0.38 which suggest either no power advantage or a minor one in favor of the CNN; this will be a factor to continue investigating and may require refinement or increasing sophistication in the ethograms derived from DeepLabCut.

The human labor associated with training the CNN and later video analysis is relatively minor. Labeling the 480 training frames took approximately 50 minutes and was done by someone with no prior training in behavioral science. Next, the CNN ran ~1 million iterations over the course of ~18 hrs on an off the shelf GPU; future applications may require fewer iterations as accuracy reached the maximum at about 500,000 cycles and performance can increase with more, or more powerful GPUs, or cloud/cluster computing. Once the CNN was trained, test videos were evaluated at approximately 2x the video rate. That is, the CNN produced x,y coordinates for 5 postures in ~½ the time needed to watch the video in real time. This rate would increase with smaller video size (i.e. lower resolution) and greater GPU power. Importantly, the CNN was able to accurately track postures in a second set of videos that were collected under similar experimental conditions but had not been sampled for training the CNN. Thus, an initial investment in labeling and model training would allow for many future applications with very little human effort. In our experience, we found that varying the bitrate, framerate, and dimensions of the source videos all influenced processing time and we encourage other adopters to experiment with these factors in their preparations.

## Appendix 1

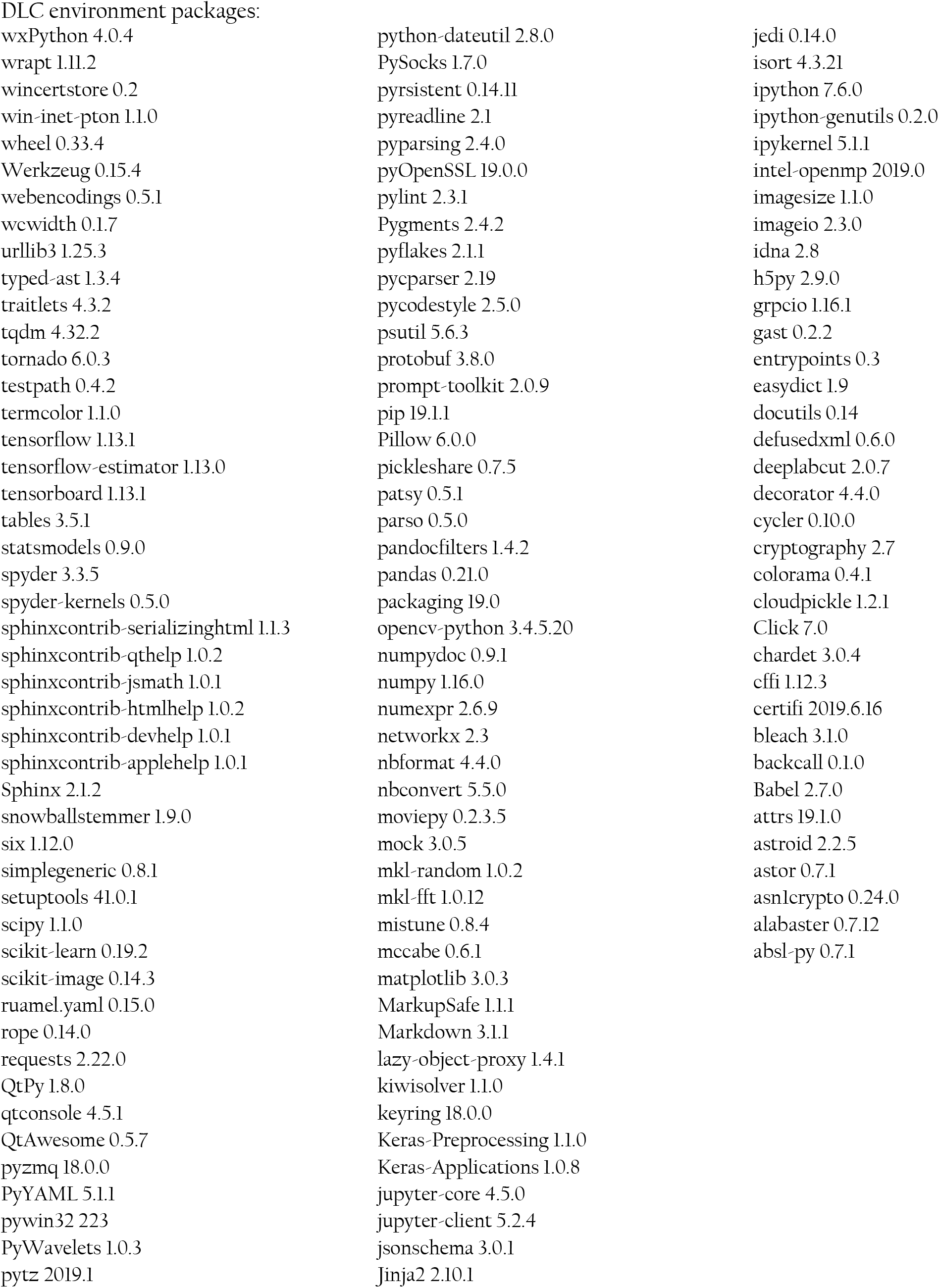

